# The clock gene *Per1* expression may exert diurnal control over hippocampal memory consolidation

**DOI:** 10.1101/2022.10.11.511798

**Authors:** Lauren Bellfy, Chad W. Smies, Alicia R. Bernhardt, Kasuni K. Bodinayake, Aswathy Sebastian, Emily M. Stuart, Destiny S. Wright, Chen-Yu Lo, Shoko Murakami, Hannah M. Boyd, Megan J. von Abo, Istvan Albert, Janine L. Kwapis

## Abstract

The circadian system influences many different biological processes, including memory performance. While the suprachiasmatic nucleus (SCN) functions as the brain’s central pacemaker, satellite clocks have also been identified in other brain regions, such as the memory-relevant dorsal hippocampus. Although it is unclear how these satellite clocks contribute to brain function, one possibility is that they may serve to exert diurnal control over local processes. Within the hippocampus, for example, the local clock may contribute to time-of-day effects on memory. Here, we used the hippocampus-dependent Object Location Memory task to determine how memory is regulated across the day/night cycle in mice. First, we systematically determined which phase of memory (acquisition, consolidation, or retrieval) is modulated across the 24h day. We found that mice show better long-term memory performance during the day than at night, an effect that was specifically attributed to diurnal changes in memory consolidation, as neither memory acquisition nor memory retrieval fluctuated across the day/night cycle. Using RNA-sequencing we identified the circadian clock gene *Period1* (*Per1*) as a key mechanism capable of supporting this diurnal fluctuation in memory consolidation, as *Per1* oscillates in tandem with memory performance. We then show that local knockdown of *Per1* within the dorsal hippocampus has no effect on either the circadian rhythm or sleep behavior, although previous work has shown this manipulation impairs memory. Thus, *Per1* may independently function within the dorsal hippocampus to regulate memory in addition to its known role in regulating the circadian rhythm within the SCN. *Per1* may therefore exert local diurnal control over memory consolidation within the dorsal hippocampus.

## Introduction

Circadian rhythms are responsible for regulating integral physiological processes across the 24-hour day in organisms big and small [1,2]. The circadian system is primarily regulated by the suprachiasmatic nucleus (SCN) in the hypothalamus [3], but satellite clocks have also been identified across the brain and body, including within memory-relevant regions like the dorsal hippocampus [4]. Although it is clear that memory performance oscillates across the day/night cycle, the mechanisms that modulate memory across the diurnal cycle are unknown. Recent work has suggested that clock genes function within memory-relevant brain regions to exert local diurnal control over memory [5–10], but it is unclear which clock genes regulate memory and which phase of memory (acquisition, consolidation, or retrieval) is impacted by the time of day.

The molecular clock begins with a CLOCK-BMAL1 heterodimer binding to E-box motifs upstream of two gene families (*Period* (*Per*) and *Cryptochrome* (*Cry*)) to induce their transcription [11]. *Per* and *Cry* are then translated in the cytoplasm and these proteins dimerize before returning to the nucleus to inhibit the CLOCK-BMAL1 complex, blocking subsequent transcription of *Per1* and *Cry* [12–14]. Proteolytic decay of PER and CRY proteins frees up the CLOCK-BMAL1 complex, which resets the feedback loop, enabling a new round of transcription of *Per1* and *Cry* [15]. This entire transcription/translation feedback loop takes ∼24 hours and is roughly aligned with the natural light/dark cycle. Notably, these core clock genes also rhythmically oscillate in most cells outside the SCN, including within neurons of memory-relevant brain structures like the dorsal hippocampus [4]. This could provide a potential mechanism through which the circadian clock modulates memory; clock genes could function within specific brain structures to exert local circadian control over memory and other region-specific functions [5,6]. In particular, previous work has shown that the core clock gene *Per1* is important for memory formation [7,9,16]. Bidirectional manipulation of *Per1* within the dorsal hippocampus modulates memory; local knockdown of *Per1* impairs spatial memory whereas local overexpression improves memory in aging mice [7]. Thus, *Per1* functions locally within the dorsal hippocampus to regulate memory, although it is not clear what phase of memory is modulated via this mechanism.

Successful long-term memory formation requires several phases: acquisition, consolidation, and retrieval. Memory is formed during the acquisition phase in which the memory is initially learned. Following acquisition, the information can be stored in either short- or long-term memory. Short-term memories, created in the absence of transcription, retain the information only transiently (typically a few hours). For long-term memory to form, transcription needs to occur around the time of learning [17–19], presumably to drive the cellular and synaptic modifications needed for long-term storage. This process of stabilizing learned information into robust and persistent long-term memory is termed consolidation. Finally, to behaviorally express the memory at a subsequent test, the memory must be properly retrieved. Long-term memory is typically tested 24h or longer after acquisition, after the consolidation process is complete, and can be very long lasting, up to the lifespan of the animal [20]. Although memory performance is clearly affected by the time of day [2,21,22], because most studies use the same diurnal timepoint for both training and testing (i.e. training and testing are separated by exactly 24h), it is unknown which phase of memory is specifically impacted across the day/night cycle.

In this study, we tested the hypothesis that memory consolidation, rather than acquisition or retrieval, is altered across the diurnal cycle. Specifically, we hypothesized that circadian clock genes function locally within the dorsal hippocampus to exert circadian control over spatial memory consolidation. To test this, first, we used Object Location Memory (OLM), to determine how hippocampus-dependent spatial memory performance in mice is affected by the time of day and found that memory is better during the day than at night. Next, we ran a series of experiments to determine which phase of memory (acquisition, consolidation, or retrieval) is specifically affected by the day/night cycle. We found that nighttime memory deficits are specifically due to altered consolidation, suggesting that learning-induced gene expression might underlie diurnal fluctuations in memory. Using unbiased RNA-seq, we then identified the clock gene *Per1* as a key mechanism that may regulate the consolidation process in a time-of-day dependent manner in the dorsal hippocampus (DH), consistent with previous work from our lab and others [7–9,16]. Finally, we show that reducing local *Per1* in the DH does not affect circadian activity patterns or sleep behavior. This demonstrates, for the first time, that *Per1* exerts local diurnal control over memory consolidation within the hippocampus in addition to its well-documented role in modulating the circadian system within the SCN.

## Materials and Methods

*Mice* Mice were young adult male C57BL/6J mice between 2 and 4 months old at time of behavioral experiments (weighing between 25-30 g). Food and water were accessible *ad libitum* and all mice were housed on a 12 h light/dark cycle, with lights turning on at 6am (7am during Daylight Saving Time). Mice trained during the day (ZT1, 5, and 9) were entrained to a standard light cycle whereas mice trained at night (ZT13, ZT17, and ZT21) were entrained to a reverse light cycle. Mice were typically group housed for acquisition, retrieval, and consolidation experiments, although a cohort of single-housed mice were used in one replication of the first experiment (Fig. 1). As all ZT groups were equally represented in this replication and no behavioral differences were observed between housing conditions in any phase of the experiment (e.g. Testing: Two-way ANOVA, significant effect of ZT Time (*F*_(5,61)_=3.896, p<0.01), no effect of Housing or Interaction), data for single-and group-housed mice were collapsed. For the circadian rhythm/sleep experiment (Fig. 5), mice were single-housed to enable accurate measurement of each individual animal’s movement. All experiments were performed according to US National Institutes of Health guidelines for animal care and use and were approved by the Institutional Animal Care and Use Committee of the Pennsylvania State University.

**Fig 1.**
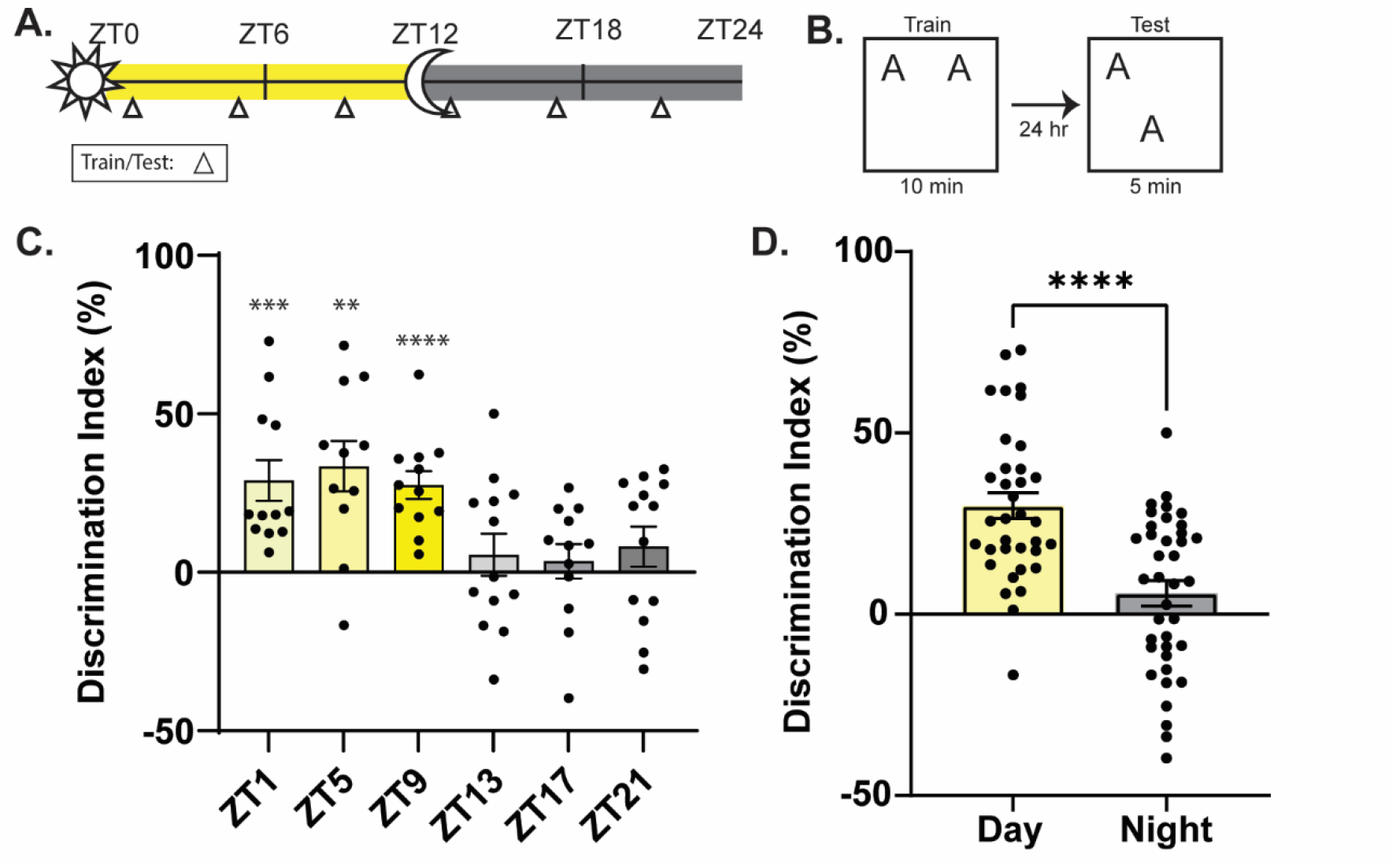
Memory performance oscillates over the diurnal cycle with better memory performance during the day. **A)** Schematic for behavioral time points. Triangle indicates time of training and testing. **B)** Object location memory experimental design. **C)** Memory performance oscillates over 24 hours with best memory observed at ZT5 and worst at ZT17. Yellow bars indicate day timepoints while grey bars indicate night timepoints (n=11-13/timepoint). **D)** Memory performance is significantly better during the day than at night (n=35-38/timepoint). ** = p<0.01, *** = p<0.001, **** = p<0.0001 compared to zero (C) or between groups (D). ZT = Zeitgeiber Time, where ZT0 = 6am (7am DST), lights on, ZT12 = 6pm (7pm DST), lights off.

*Object Location Memory (OLM)* OLM was conducted as previously described [7,23]. Mice were handled for 2 min/day for 4 days and habituated to the context for 5 min/day for 6 days. For training, mice were exposed to two identical objects (100 mL beakers filled with cement) for 10 minutes. For testing, mice were returned to the context for 5 minutes with one object moved to a new location. For long-term memory tests, mice were placed back into the context 24 or 36 hours later and for short-term memory tests, mice were placed back into the context 60 minutes later. Object exploration was quantified by blinded experimenters and exploration was defined as the mouse orienting to the center of the object and within 1 cm. Total time spent investigating was quantified with DeepEthogram [24], with the scoring parameters chosen to closely match handscoring methods previously used to score OLM behavior [7,23,25]. A discrimination index was calculated based off the amount of time spent investigating the novel location in comparison to the original location. Mice that explored <3s during testing, <5s during training, or that showed an existing object preference (DI > ±20) during training were removed from all analyses. Habituation sessions were analyzed for distance traveled and speed using Ethovision (Noldus). For molecular experiments, mice were sacrificed via cervical dislocation 60 minutes following training and the brain was harvested and flash-frozen in isopentane.

*RT-qPCR* Tissue was harvested from DH punches and frozen at −80°C until use. RNA was extracted using the RNeasy Minikit (Qiagen). cDNA was synthesized using the Hi-Fi cDNA Synthesis Kit (Abcam). To reduce the potential for genomic DNA amplification, assays were designed to be intron spanning for the *Per1* target assay and the *GAPDH* reference assay (IDT) designed based on the Mus_musculus, GRCm38 build. qPCR was run on the Roche LightCycler with the following cycling conditions: 1 cycle at 95°C for 3 min, 45 cycles of 95°C for 15 sec and 60°C for 60 sec, and hold at 4°C. Data were analyzed with LightCycler Analysis software using the Relative Quantification analysis method. For *Per1* we used the following primers: left 5’-CCTGGAGGAATTGGAGCATATC-3’; right 5’-CCTGCCTGCTCCGAAATATAG-3’; probe 5’-AAACCAGGACACCTTCTCTGTGGC-3’ labeled with FAM. We used *Gapdh* as a reference assay with the following primers: left 5’-GGAGAAACCTGCCAAGTATGA-3’; right 5’-TCCTCAGTGTAGCCCAAGA-3’; probe 5’-TCAAGAAGGTGGTGAAGCAGGCAT-3’ labeled with HEX.

*RNA-Sequencing* RNA was extracted from dorsal hippocampus (DH) 500 µm punches and RNA quality and concentration was evaluated using RNA 6000 pico kit for Bioanalyzer (Agilent) with a cutoff of 9 for the RIN. Data consisted of 8-53 bp paired end reads. FastQC reports were generated to check the data quality. Data were mapped to the reference genome (Mus_musculus, GRCm38 build) using hisat2 (v2.1.0). The average mapping rate was 95%. Mapped data was visualized and inspected in IGV (Integrative Genomics Viewer). Reads that mapped to the genes were counted using featureCounts (v2.0.0) specifying -s 2 -p parameters. Transcript level counts were obtained using Kallisto (v0.46.1). Raw reads were normalized and differentially expressed genes and transcripts were obtained using DESeq2 package. Circadian genes were identified with JTK algorithm specifying 24h window in Metacyle program. Pathway analysis were preformed using KEGG database. Over-representation analysis with clusterProfiler package in R was used to identify enriched KEGG pathways.

*HSV Production* Neuron-specific herpes simplex viruses (HSVs) were used to express the CRISPR inhibition (CRISPRi) system (dCas9-KRAB-MeCP2) and the single guide RNA (sgRNA), either targeting *Per1* or a non-targeting control. All viruses were purchased from Dr. Rachael Neve (Gene Delivery Technology Core, Massachusetts General Hospital, Boston, MA). CRISPRi was expressed in a p1005 vector that simultaneously expresses a deactivated Cas9 (dCas9)-KRAB-MeCP2 fusion under the hSyn promoter along with mCherry (via an IRES element). The *Per1* sgRNA was previously designed and chosen based on its efficacy in reducing *Per1* in HT22 cell culture and has previously been shown to locally reduce *Per1* in the brain *in vivo* [8].

The *Per1* sgRNA or the non-targeting control sgRNA was cloned in the pDonr221 vector (ThermoFisher) containing the U6 promoter and the sgRNA sequence. Dr. Neve subcloned the inserts into an HSV vector with GFP under the CMV promoter. The sgRNA sequences were as follows: *Per1*: GAGTTCGACGGCTCCAGAGTA and non-targeting: GCGAGGTATTCGGCTCCGCG. The non-targeting sgRNA was previously validated in similar CRISPR systems [26].

*Stereotaxic Surgery* Mice were anesthetized with 1.5%-4% isoflurane and placed in the stereotaxic apparatus. After leveling the skull, Bregma was measured and holes were drilled through the skull above the dorsal hippocampus (DH; AP, −2.0 mm; ML, ± 1.5 mm, DV, −1.5 mm relative to Bregma). Bilateral injection needles were lowered to the DH at a rate of 0.2 mm/15 s. Two minutes after reaching the DH, 2.0 µL of an equal parts mixture of HSV-CRISPRi and HSV-sgRNA-*Per1* or HSV-sgRNA-ctrl was infused into each hemisphere at a rate of 10 µL/h. Post-injection, injection needles remained in place for 5 minutes, then moved up 0.1 mm and remained in place for an additional 5 minutes prior to being removed at a rate of 0.1 mm/15 s.

*Circadian Rhythm and Sleep Time Analysis* Mice were single housed under a 12 h light/dark (LD) cycle and their activity was continuously monitored with passive infrared sensors and Clocklab software (both Actimetrics). After a 2-week entrainment period, mice were infused with HSV-CRISPRi and either the HSV-sgRNA-*Per1* or HSV-sgRNA-ctrl (described above) to reduce *Per1* in the dorsal hippocampus. Mice were given 2 days of recovery to enable the virus to fully express before being placed in constant darkness (DD) for 10 days (after which the HSV is fully eliminated [27,28]). Tau values were calculated using the onset of free activity and calculating the least-squares fit with Clocklab software. Data obtained between surgery and constant darkness was not included in the analysis, as mice were removed from their homecages and weighed daily during this recovery period, rendering the movement/sleep data inaccurate. Sleep behavior was assessed using a program adapted from the COMPASS system [29].

Movement data were collected in 10s bins with the criteria that periods of inactivity lasting 40s or longer were quantified as a behavioral correlate of sleep. Previous work has demonstrated a high degree of correlation between this immobility-defined sleep and sleep defined using EEG records [29]. This sleep behavior (in seconds) was combined within 30 minute bins across the day/night cycle.

*Statistical Analysis* Differences in memory performance were analyzed using one sample *t*-tests (Fig. 1C, to compare each group’s object preference to zero (no preference)), unpaired *t*-tests, or two-way ANOVAs followed by Sidak’s multiple comparison *post-hoc* tests. For RT-qPCR, each group was normalized to the ZT1 homecage. Sleep behavior, in which each animal’s sleep was measured multiple times across the day/night cycle, was analyzed with mixed-model ANOVAs, in which sleep behavior over time was analyzed as a repeated measures variable. For all analyses, significance was indicated by an α value of 0.05.

## Results

### Memory performance oscillates over the diurnal cycle

Memory performance is known to oscillate over the 24-hour day. To identify when spatial memory is best and worse across the day/night cycle, mice were trained in dorsal hippocampus-dependent object location memory (OLM) at 6 distinct Zeitgeber Times (ZTs): ZT1, ZT5, ZT9, ZT13, ZT17, and ZT21, where ZT0 = lights on and ZT12 = lights off (Fig. 1A). Mice were tested 24 hours after training to specifically assess their long-term memory (LTM) performance at the same diurnal timepoint (Fig. 1B).

Consistent with previous reports [2,8,16,30,31], we found that memory was best during the day and worst at night (Fig. 1C-D). Specifically, we found that mice showed robust memory at all of the day timepoints, ZT1, ZT5, and ZT9, (One-sample *t*-test compared to 0, ZT1: *t*_(11)_=4.527, p<0.001, ZT5: *t*_(10)_=4.214, p<0.01, ZT9: *t*_(11)_=6.336, p<0.0001), and poor memory during the night, as indicated by a DI near 0, indicating no preference for the moved object (ZT13: *t*_(12)_=0.8414, p=1.4166, ZT17:*t*_(11)_=0.6394, p=0.5356), although memory performance began to improve as the night is ending at ZT21 (*t*_(12)_=1.290, p=0.2214). Memory peaked at ZT5, in the middle of the light cycle, and showed a trough at ZT17, in the middle of the dark cycle (Fig. 1C). When all of the day and night timepoints were collapsed, we found that memory was significantly better during the day compared to at night (Fig 1D; Unpaired *t*-test, *t*_(71)_=4.840, p<0.0001). Therefore, memory performance oscillates over the day/night cycle, with better memory performance occurring during the day and worse memory at night.

### OLM acquisition is intact across the day/night cycle

After confirming that memory oscillates over the 24-hour day, we next wanted to determine which phase of memory is specifically regulated by the day/night cycle: acquisition, consolidation, or retrieval. First, to test whether acquisition changes across the 24h day, we measured short-term memory (STM) for OLM during the day and night. We trained a cohort of mice in OLM at the peak of memory, ZT5, or the trough of memory, ZT17 (Fig. 2A), as identified in the previous experiment (Fig. 1C). Here, however, mice were tested 60 minutes after training (Fig. 2B) during the transcription-independent STM phase [17,32]. We found that all mice showed intact short-term memory regardless of the time of acquisition (Fig. 2C; Unpaired *t-*test comparing ZT5 to ZT17: *t*_(12)_=0.3749, p>0.05; One-sample *t*-test comparing each group to 0: ZT5: *t*_(6)_=4.801, p<0.01; ZT17: *t*_(6)_=11.77, p<0.0001). Even mice trained at ZT17 that show drastic impairments in long-term memory (Fig. 1C) showed intact memory when tested 60m after acquisition. This demonstrates that mice successfully learn OLM at night, even when long-term memory fails. Therefore, acquisition is not affected by the time of day and the observed nighttime deficits in long-term memory are not due to an acquisition deficit but are more likely due to a consolidation or retrieval deficit.

**Fig 2.**
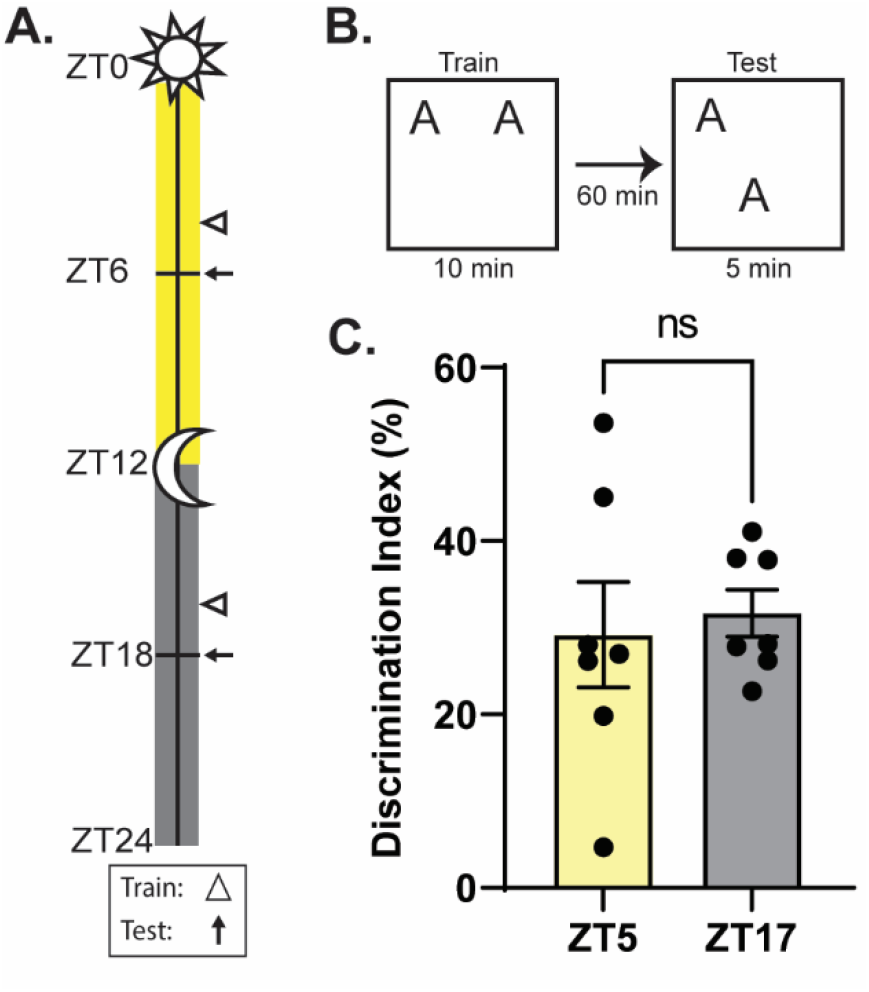
Short term memory is intact during the day and the night. **A)** Schematic for behavioral time points. Triangle indicates time of training. **B)** Object location memory experimental design. **C)** No difference is seen in short term memory performance between the day (yellow bar) and the night (grey bar; n=7/timepoint). ns = not significant, ZT = Zeitgeiber Time, where ZT0 = 6am (7am DST), lights on, ZT12 = 6pm (7pm DST), lights off.

### OLM retrieval is also stable across the day/night cycle

After ruling out acquisition as driving time-of-day effects on memory performance, we next wanted to test whether memory retrieval is altered across the day/night cycle. It is possible that mice tested at night have intact memory for OLM but have difficulty retrieving that stored information during the dark cycle. To rule out a retrieval deficit, we again trained animals at the peak and trough of memory (Fig. 3A) and tested them either 24 hours (at the same ZT) or 36 hours later (at the opposite ZT) (Fig. 3B) to separate the acquisition ZT from the retrieval ZT. We found that the time of acquisition, and not the time of retrieval, drove memory performance in OLM. Regardless of when they were tested, mice trained during the daytime (ZT5) showed strong memory (Fig. 3C) whereas mice trained at night (ZT17) showed weak object location memory (Fig. 3C). We saw a significant difference in memory performance between the groups trained during the day compared to those trained at night (Two-way ANOVA, significant effect of Training Time (*F*_(1,22)_=43.57, p<0.0001), no effect of Retrieval Time or Interaction), but no significant difference within the day-trained cohorts or the night-trained cohorts (Sidak’s *post-hoc*, p>0.05). Thus, if memory acquisition occurred during the day, memory was successfully retrieved at test and if memory acquisition occurred at night, retrieval was impaired at both day and night timepoints, suggesting that retrieval itself is not altered across the day/night cycle. Together with the results of Figure 2, our work suggests that hippocampal memory consolidation, rather than memory acquisition or retrieval, oscillates across the 24h day.

**Fig 3.**
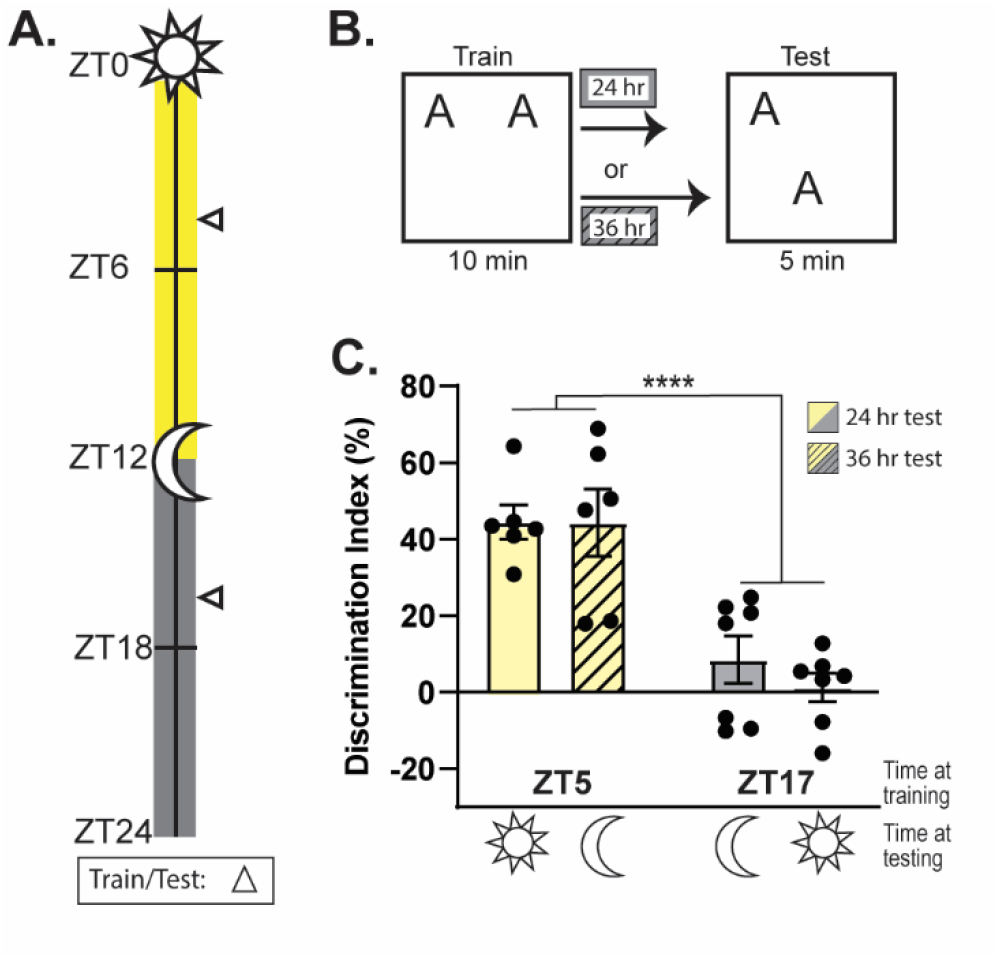
Memory performance is determined by time of consolidation not time of retrieval. **A)** Schematic for behavioral time points. Triangle indicates time of training and/or testing. **B)** Object location memory experimental design. Testing was performed 24h (solid bars) or 36h (striped bars) post-training. **C)** Mice trained during the day (yellow bars) showed strong memory performance whether tested 24h (day) or 36h (night) later. Mice trained at night (grey bars) showed weak memory performance whether tested 24h (night) or 36h (day) later (n=6-7/timepoint). **** = p<0.0001, ZT = Zeitgeiber Time, where ZT0 = 6am (7am DST), lights on, ZT12 = 6pm (7pm DST), lights off.

### Daytime learning drives major changes in gene expression

Our behavioral findings suggest that the observed nighttime memory deficits are specifically due to a consolidation error, as both memory acquisition and memory retrieval were intact at night. As *de novo* transcription is critically important for the memory consolidation process [18,20,33], we reasoned that changes in learning-induced gene expression could underlie these nighttime memory deficits. Specifically, we hypothesized that a subset of learning-induced genes that support memory during the daytime are not properly expressed at night, leading to the observed impairments in long-term memory consolidation. Although previous work has suggested that clock genes (notably *Per1*) function within the hippocampus to regulate memory, we wanted to identify all potential genes capable of exerting diurnal control over hippocampal memory consolidation in an unbiased manner. We therefore used RNA sequencing (RNA-seq) to identify genes that oscillate in tandem with memory consolidation across the day/night cycle. To this end we trained mice in OLM at 6 timepoints across the diurnal cycle, as in Fig. 1A and then sacrificed them 60 minutes later (along with time-matched homecage controls) to assess RNA expression in the DH (Fig. 4A). After collecting punches from the dorsal hippocampus (area CA1) and isolating RNA, we then created libraries and ran RNA-seq to identify genes capable of enabling robust memory during the day but poor memory at night.

**Fig 4.**
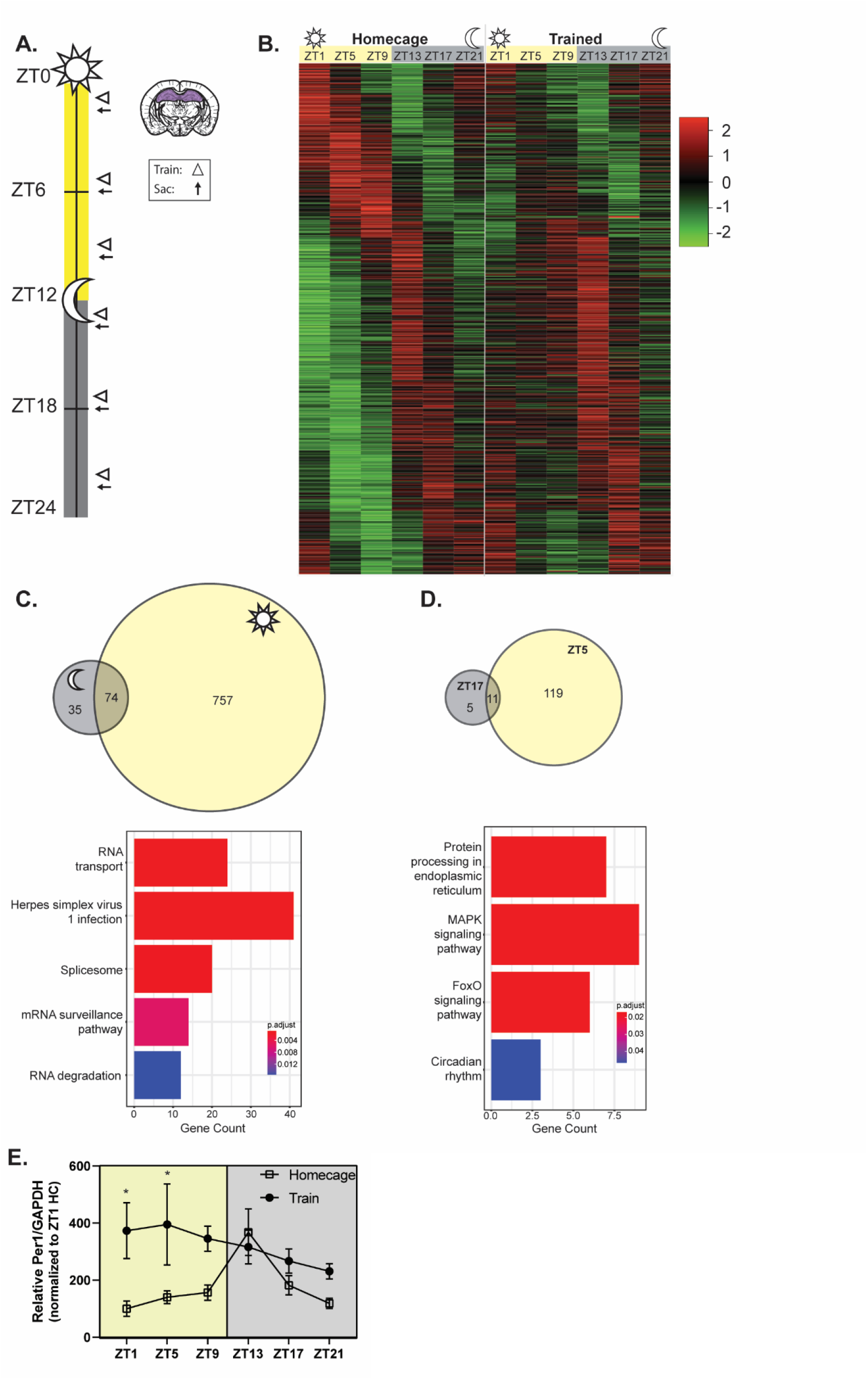
Learning during the day drives major changes in gene expression that do not occur at night. **A)** Schematic for behavioral and sacrificing timepoints. Triangle: training time; arrow: sacrifice time. **B)** Oscillatory genes in homecage mice (left) show drastic learning-induced changes during the day (right, yellow columns) but not at night (gray columns). **C)** Top: genes upregulated in response to learning during the day (yellow), night (gray) or both. Bottom: top Kegg pathways upregulated during the day but not at night. **D)** Top: genes upregulated in response to learning at ZT5 (yellow), ZT17 (gray), or both. Bottom: top Kegg pathways upregulated at ZT5 but not at ZT5 but not at ZT17. **E)** *Per1* mRNA is increased by learning during the day (ZT1 and ZT5) but not at any night timepoint (n=4-8/timepoint). * = p<0 05 ZT = Zeitgeiber Time, where ZT0 = 6am (7am DST), lights on, ZT12 = 6pm (7pm DST), lights off.

First, we used JTK-Cycle to identify genes that oscillate across the day/night cycle [34]. To see how learning changes oscillatory gene expression, we first identified genes that oscillate under HC conditions (left column) and then plotted the same genes for the OLM trained group (right column). We found that during the day (ZT1, ZT5, ZT9), OLM drives dramatic changes in gene expression whereas during the night (ZT13, ZT17, ZT21) fewer genes were affected by the same training event, with most genes continuing to oscillate normally even after OLM (Fig. 4B). This shows that learning is massively disruptive to baseline gene oscillations during the day, but not at night. This is consistent with our hypothesis that a subset of learning-induced genes fail to respond to learning at night, potentially contributing to nighttime reductions in memory consolidation.

Next, we aimed to identify individual genes capable of modulating memory across the day/night cycle. A number of genes are known to be induced by learning and necessary for memory consolidation [18,33,35], but there is very little known about how these learning-induced genes oscillate across the 24h day. As we specifically wanted to identify genes that might exert circadian control over memory (supporting robust memory during the day but only weak memory at night), we aimed to identify genes that show learning-induced increases during the daytime that are reduced or eliminated at night in tandem with memory performance (Fig. 2). We therefore used differential gene expression analyses to identify genes induced by learning (i.e. genes expressed at significantly higher levels in mice trained with OLM compared to time-matched homecage controls) at each ZT and then compared these learning-induced genes across the day and night to specifically identify genes capable of supporting robust learning during the day but not at night.

First, we compared genes induced by OLM during the day (at ZT1, 5, or 7) to those induced by OLM during the night (ZT13, 17, or 21). We identified 757 genes upregulated only during the day, 35 genes upregulated only at night, and 74 that were upregulated by learning during both the day and the night (Fig. 4C). This is consistent with our JTK-cycle results, indicating that OLM training drives massive changes in gene expression during the daytime (when memory is robust) that are muted at night (when memory is weak). Next, to understand the functional relevance of the genes that might support diurnal oscillations in memory, we ran pathway analyses on genes exclusively upregulated during the day using the KEGG database. During the daytime, when memory is robust, the most highly upregulated pathways dealt with RNA processing, transport, and degradation (Fig. 4C). This finding was unsurprising as *de novo* gene expression is critical to long-term memory formation [18,20,33].

To narrow down this list, we decided to restrict our analyses to directly compare learning-induced genes when memory is best (ZT5) and worst (ZT17). We identified 119 genes upregulated in response to learning exclusively at ZT5, 5 upregulated only at ZT17, and 11 upregulated at both times (Fig. 4D). Again, to determine the functional identity of genes capable of supporting diurnal oscillations in memory, we ran Kegg pathway analyses on genes that were upregulated exclusively at ZT5. The top pathways identified genes involved in protein processing in endoplasmic reticulum, MAPK signaling, FoxO signaling, and the circadian rhythm (Fig. 4D). We were particularly intrigued to see that circadian rhythm genes were induced by learning during the day but not at night, as this would suggest that clock genes might function locally in the hippocampus to exert circadian control over memory, as previously hypothesized [5,6,36]. Of these, one gene in particular stood out: *Period1* (*Per1*). *Per1* has a well-established role in establishing the circadian rhythm within the brain’s central pacemaker, the suprachiasmatic nucleus [12–14], but has more recently been implicated in learning and memory as well [7–10,16]. *Per1* has specifically been hypothesized to play a role in “gating” memory formation across the diurnal cycle [36], although the precise role that local *Per1* plays in the dorsal hippocampus is unclear. In addition to *Per1*, there were two other circadian rhythm genes differentially regulated in response to learning during the day but not the night: *Protein Kinase AMP-Activated Non-Catalytic Subunit Beta 1* (*Prkab1*) and *Basic Helix-Loop-Helix Family Member E40* (*Bhlhe40*). The roles of these genes in learning have yet to be explored.

To get a better understanding of how *Per1* oscillates across the day/night cycle within the dorsal hippocampus, we ran RT-qPCR on these samples to measure *Per1* mRNA across the day/night cycle in both homecage and trained mice. In homecage controls, we observed rhythmic oscillations in hippocampal *Per1* that peaked at the beginning of the night ((1-way ANOVA just on HC group): 1-way ANOVA, *F*_(5,39)_=5.321, *p*<0.001, Sidak’s *post hoc* test comparing ZT1 to ZT13,\*\*\**p*<0.001; no other timepoints different). Following learning, *Per1* was induced by learning during the daytime but this induction was dampened at night. We found a significant increase in *Per1* response to OLM (2-way ANOVA, significant effect of Training (*F*_(1,69)_=15.58, p=0.0002), but no effect of ZT Time or Interaction). Sidak’s *post hoc* tests comparing homecage and trained groups within each timepoint revealed that *Per1* was significantly upregulated by OLM at ZT1 and ZT5 (p<0.05) but not at any other timepoint (Fig. 4E; Sidak’s *post-hoc*, p>0.05). Thus, overall, hippocampal *Per1* is induced by learning during the day, but this induction largely fails at night, as indicated by our RNA-seq (Fig. 4D). Together with our behavioral data, this demonstrates that *Per1* oscillates in tandem with spatial memory consolidation; both memory performance and hippocampal *Per1* peak during the daytime and trough at night. *Per1* may therefore be capable of exerting local circadian control over hippocampal memory, with nighttime reductions in learning-induced *Per1* limiting memory formation.

### Manipulation of *Per1* expression in the dorsal hippocampus does not affect circadian rhythmicity or sleep patterns

Given that *Per1* plays a key role in the central circadian system, which itself can modulate memory [2,30,37–42], we wanted to ensure that our local knock-down of *Per1* in the DH does not indirectly affect memory by disrupting the central clock in the SCN. Specifically, we wanted to ensure that hippocampal *Per1* knockdown has no effect on either the circadian rhythm or sleep behavior. To knock down *Per1* we used a CRISPR inhibition (CRISPRi) system, which consists of a dead Cas9 (dCas9) fused to two transcriptional repressors: KRAB and MeCP2 (Fig. 5A) packaged in an HSV to drive neuron-specific knockdown of *Per1* [43]. HSV-CRISPRi was injected directly into the CA1 region of the DH along with either *Per1* sgRNA or non-targeting control sgRNA.

**Fig 5.**
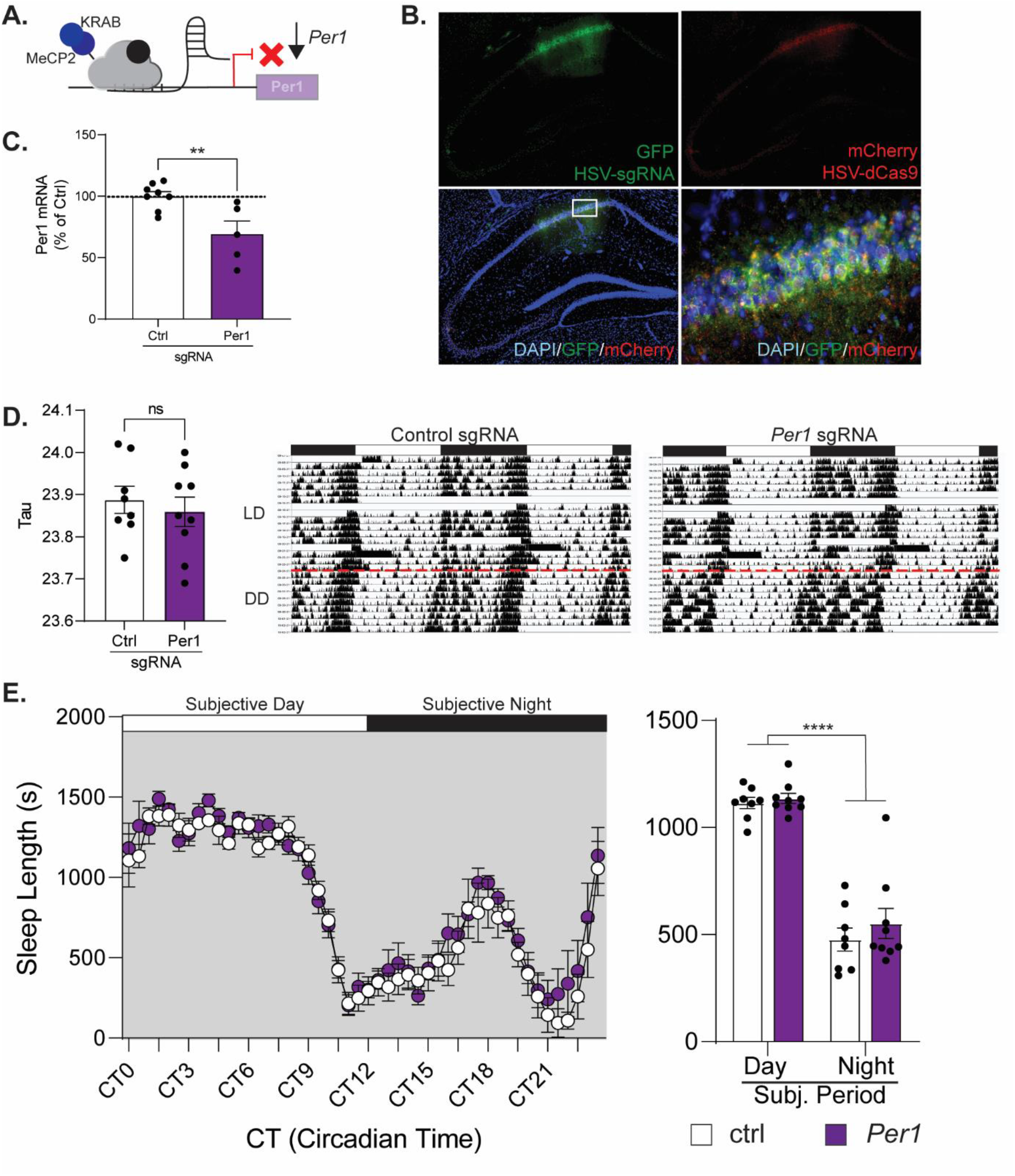
Knocking down *Per1* in the dorsal hippocampus does not affect circadian rhythms or sleep behavior. **A)** Schematic of CRISPRi system. dCas9 is fused with two repressive elements that reduce the transcription of endogenous *Per1*. **B**) Representative HSV-CRiSPRi expression in the DH 3d post-injection, with a high degree of co-expression of the sgRNA vector (GFP, top left) and dCas9 (mCherry, top right). Bottom: merged images. **C)** HSV-CRISPRi significantly reduces *Per1* expression *in vivo* (n=5-8/condition). **D)** Knocking down *Per1* in the DH does not affect free-running tau under dark-dark conditions. Right: representative actograms for control sgRNA (middle panel) and *Per1* sgRNA (right panel). **E)** Sleep length was not affected by Pen knockdown. Right: average sleep length for the subjective day and night Mice showed longer sleep bouts during the subjective day, as expected, but there was no effect of *Per1* knockdown (n=8-9/condition). **= p<0 01, **** = p <0.0001, ns = not significant, LD = light/dark, DD = dark/dark. CT = Circadian Time. ZT = Zeitgeiber Time. where ZT0 = 6am (7am DST), lights on, ZT12 = 6pm (7pm DST), lights off.

First, to confirm that HSV-CRISPRi significantly reduces hippocampal *Per1* expression, a subset of mice was sacrificed three days after injection (when HSV expression peaks) for immunofluorescence and qPCR. We observed high colocalization of the sgRNA (green) and dCas9-KRAB-MeCP2 (red) in neurons of the dorsal hippocampus (Fig. 5B). Further, in punches collected from this region, we found that *Per1*-targeting CRISPRi significantly decreases the expression of *Per1* in the DH compared to controls, with a decrease of 30.84% (Fig. 5C; Unpaired t-test *t*_(11)_=3.252, p<0.01). Therefore, our HSV-CRISPRi system appropriately reduces *Per1* expression within the DH.

To test whether hippocampal *Per1* knockdown affects the animals’ circadian activity pattern, a separate cohort of mice underwent activity monitoring in LD conditions followed by DD conditions. Mice were acclimated to the standard LD cycle for 2 weeks prior to injection of CRISPRi into the DH. Three days post-injection, at the peak of viral expression, the lights were turned off and activity and sleep behavior were monitored in constant darkness for 10 days (for the duration of HSV expression). We found that local *Per1* knockdown within the dorsal hippocampus had no significant effect on circadian activity patterns. There was no significant difference in free-running τ between groups (*Per1* knockdown mice: 23.86, control mice: 23.91 (Fig. 5D; Unpaired *t*-test *t*_(15)_=0.5955, p=0.5604) indicating that the circadian rhythm was intact even in mice with hippocampal *Per1* knockdown. Therefore, hippocampus-specific knockdown of *Per1* does not affect the circadian activity pattern.

We also assessed sleep behavior in these mice. Briefly, we used our infrared monitors to identify bouts of inactivity lasting 40 sec or longer as a behavioral correlate of sleep. This immobility-defined sleep has previously been shown to tightly correlate with sleep defined via EEG records [29]. We observed no differences in the sleep behavior (including both sleep duration and sleep bout length) between *Per1* knockdown and control mice in either the LD or the DD phase (Fig. 5E; LD: Two-way mixed-model ANOVA, effect of Time (*F*_(1,15)_=289.6, p<0.0001), no effect of Injection or Time x Injection). Therefore, knocking down *Per1* in the DH does not affect sleep behavior (Fig. 5E) or circadian activity patterns (Fig. 5D). Along with previous research showing that hippocampus-specific manipulations of HDAC3 (a major epigenetic regulator of *Per1*) [7] and even electrolytic lesions of the DH [44] have no effect on the circadian rhythm, this work strongly suggests that the sleep/wake cycle is not affected by site-specific manipulations in the dorsal hippocampus.

## Discussion

Memory is carefully regulated by the circadian system, but the mechanisms that control memory across the day/night cycle are largely unclear. Here, we show that hippocampal memory oscillates across the diurnal cycle, with memory peaking during the daytime (specifically at ZT5) and showing a trough at night (ZT17). Next, through a series of experiments, we determined that memory consolidation, not memory acquisition or retrieval, is impacted by the time of day. Using RNA-seq, we next determined that learning drastically affects oscillating gene patterns specifically during the daytime and identified the circadian gene *Per1* as a key player potentially capable of exerting diurnal control over memory. Finally, we verified that *Per1* manipulations restricted to the dorsal hippocampus have no effect on either circadian activity patterns or sleep behavior. Together, these data suggest that *Per1* may play a local, autonomous role in the dorsal hippocampus to exert diurnal control over memory consolidation in addition to its well-documented role in regulating the circadian system within the SCN.

Our work suggests that hippocampal *Per1* could regulate memory based on the time of day, a modulatory role that may not be specific to the hippocampus. Our lab has recently shown that *Per1* levels increase in response to learning in another memory-relevant structure, the anterior retrosplenial cortex (aRSC) [8]. As in this study, local knockdown of *Per1* within the aRSC before learning (in this case, context fear conditioning) impaired memory, indicating *Per1* modulates multiple forms of memory consolidation across different memory-relevant brain regions. Interestingly, our previous work found that retrosplenial *Per1* may modulate memory in a sex-specific manner [8], with overexpression of *Per1* having different effects in male and female mice. Here, to achieve the necessary power (particularly in circadian experiments with 6-12 groups), we used only male mice. We are currently investigating these effects in female mice in a parallel set of experiments.

In our systematic and controlled diurnal memory experiment, we found that mice showed better memory performance during the day than at night. Other groups have identified similar diurnal memory patterns [2,8,16,30,31], but some studies have shown that mice have better memory at night in some tasks [45–47]. Although it is not clear why this variability exists, it may be due to differences in either the memory task or experimental procedure. For example, memory tasks that require the participation of other brain structures with different oscillatory patterns might change when memory is best and worst. Further, procedural differences, such as the use of overhead lights during the day but dim red lighting at night might affect the peak performance of the animals. Here, we carefully controlled the conditions to be able to directly compare performance in hippocampus-dependent OLM across the day/night cycle and found that memory was much better during the day than at night. This was somewhat surprising, as mice are nocturnal, but many species similarly perform better during the day than at night regardless of their active time [2]. This suggests that a species’ diurnal activity pattern is not a reliable predictor of memory performance across the 24h day. A better predictor of memory performance might be something happening at the cellular or molecular level, like local *Per1* induction or even the spontaneous activity of SCN cells, which are more responsive during the day in both nocturnal and diurnal animals [48]. Future work should therefore systematically determine whether other forms of memory, including those that do not require the hippocampus, show a similar oscillation, peaking during the day.

Our experiments demonstrate that memory consolidation is specifically modulated across the diurnal cycle, as short-term memory is intact even at night (Fig. 2C) and memory retrieval itself did not oscillate across the day/night cycle (Fig. 3C). Notably, in our retrieval experiment (Fig. 3), both cohorts tested at 36h had a full sleep cycle between acquiring the memory and retrieving it, but only the daytime-trained mice were able to successfully remember the object locations. Further, these results also suggest that the training event does not simply serve as a zeitgeber that selectively improves memory at that specific timepoint; mice trained during the daytime showed good memory even when tested 36h later in the middle of the night. This, along with the observation that memory is better during the day (when mice are normally asleep) suggests that our memory effects do not occur simply because the behavioral task disrupts the animals’ sleep.

Together, our work suggests that *Per1* plays two key roles in the brain: the canonical role of regulating the circadian clock within the SCN and a noncanonical role in exerting diurnal control over memory consolidation in the DH. We have previously identified *Per1* as an important mechanism that contributes to age-related hippocampal memory impairments in aging, 18-month-old mice [7]. Here, we show that *Per1* may specifically function within the dorsal hippocampus to exert local circadian control over memory consolidation, independent of its canonical role in regulating the circadian system within the SCN.

## Acknowledgements

This research was funded by NIH grants R01AG074041 (J.L.K.), K99/R00AG056586 (J.L.K.) and R21AG068444 (J.L.K.), Whitehall Foundation Grant #2020-05-06 (J.L.K.), American Federation for Aging Research Grant #A21105 (J.L.K), startup funds from the Eberly College of Science and Department of Biology at Pennsylvania State University (J.L.K.) and the National Institute on Aging under Grant T32 AG049676 to The Pennsylvania State University (L.B.). We would like to thank Dr. Rachael Neve and the Gene Delivery Technology Core at Massachusetts General Hospital for help designing and packaging all HSV viruses described here. We would also like to thank the Huck Institutes of the Life Sciences at Penn State for funding and assistance with RNA-sequencing.

## Notes

### Competing Interest Statement

The authors have declared no competing interest.

